# Antibody evasion and receptor binding of SARS-CoV-2 RW.1.1

**DOI:** 10.64898/2026.09.10.750645

**Authors:** Kristin Daniel, Hsiang Hong, Chien-Yu Huang, Aubree Gordon, Yicheng Guo, David D. Ho, Ian A. Mellis

**Affiliations:** Aaron Diamond AIDS Research Center, Columbia University Vagelos College of Physicians and Surgeons, New York, NY, USA; Department of Pathology and Cell Biology, Columbia University Vagelos College of Physicians and Surgeons, New York, NY, USA; Department of Microbiology, National Taiwan University College of Medicine, Taipei, Taiwan; Department of Epidemiology, University of Michigan, Ann Arbor, MI, United States of America; Division of Infectious Diseases, Department of Medicine, Columbia University Vagelos College of Physicians and Surgeons, New York, NY, USA; Department of Microbiology and Immunology, Columbia University Vagelos College of Physicians and Surgeons, New York, NY, USA; Pandemic Research Alliance unit at the Wu Center for Pandemic Research, Columbia University Vagelos College of Physicians and Surgeons, New York, NY, USA

## Abstract

The continued evolution of SARS-CoV-2 is shaped by changes in antibody evasion and receptor engagement that influence viral fitness. RW.1.1, an emerging descendant of XFJ carrying five additional spike substitutions, has recently increased in frequency in North America. Here, we characterized the serum antibody evasion, monoclonal antibody sensitivity, and ACE2 receptor engagement of RW.1.1. Pseudovirus neutralization assays using sera from adults showed that RW.1.1 was not more resistant to serum neutralization than currently circulating variants, with neutralizing titers comparable to XFG, BA.3.2.2, and NB.1.8.1 in the tested cohort. Neutralization of RW.1.1 also did not differ significantly among adults, children, and infants and toddlers. Despite the absence of increased overall serum antibody resistance, monoclonal antibody neutralization assays revealed substantial resistance to several RBD class 1 antibodies and increased resistance to a subset of class 1/4 antibodies. Moreover, RW.1.1 exhibited reduced ACE2 receptor engagement compared with XFG, which itself has reduced receptor engagement relative to earlier JN.1 subvariants. Thus, RW.1.1 has continued to expand despite a further reduction in receptor engagement and without a substantial increase in overall serum antibody evasion. These findings suggest that the fitness of emerging SARS-CoV-2 variants may depend not only on the magnitude of antibody evasion but also on the specific components of the polyclonal antibody response that are evaded, highlighting the increasingly complex interplay between population immunity and receptor engagement in shaping SARS-CoV-2 evolution.

## Main Text

The continued evolution of SARS-CoV-2 has generated successive variants with changes in susceptibility to antibody-mediated neutralization and receptor engagement that contribute to their growth advantage, necessitating the periodic reassessment of vaccine composition, monoclonal antibody authorizations, and other public health interventions^1^. Recently, the recombinant JN.1-derived variant XFG outcompeted NB.1.8.1 in North America despite exhibiting reduced ACE2 receptor affinity, coincident with increased antibody evasion^2^. RW.1.1, an emerging descendant of the recombinant JN.1 subvariant XFJ carrying five additional spike substitutions (**Figure 1A**), has subsequently increased in frequency in North America, accounting for approximately 10% of sequenced infections since May 2026 (**Figure S1**). We therefore investigated whether the emergence of RW.1.1 was associated with further changes in serum antibody evasion, epitope-specific monoclonal antibody resistance, or ACE2 receptor engagement.

**Figure 1.**
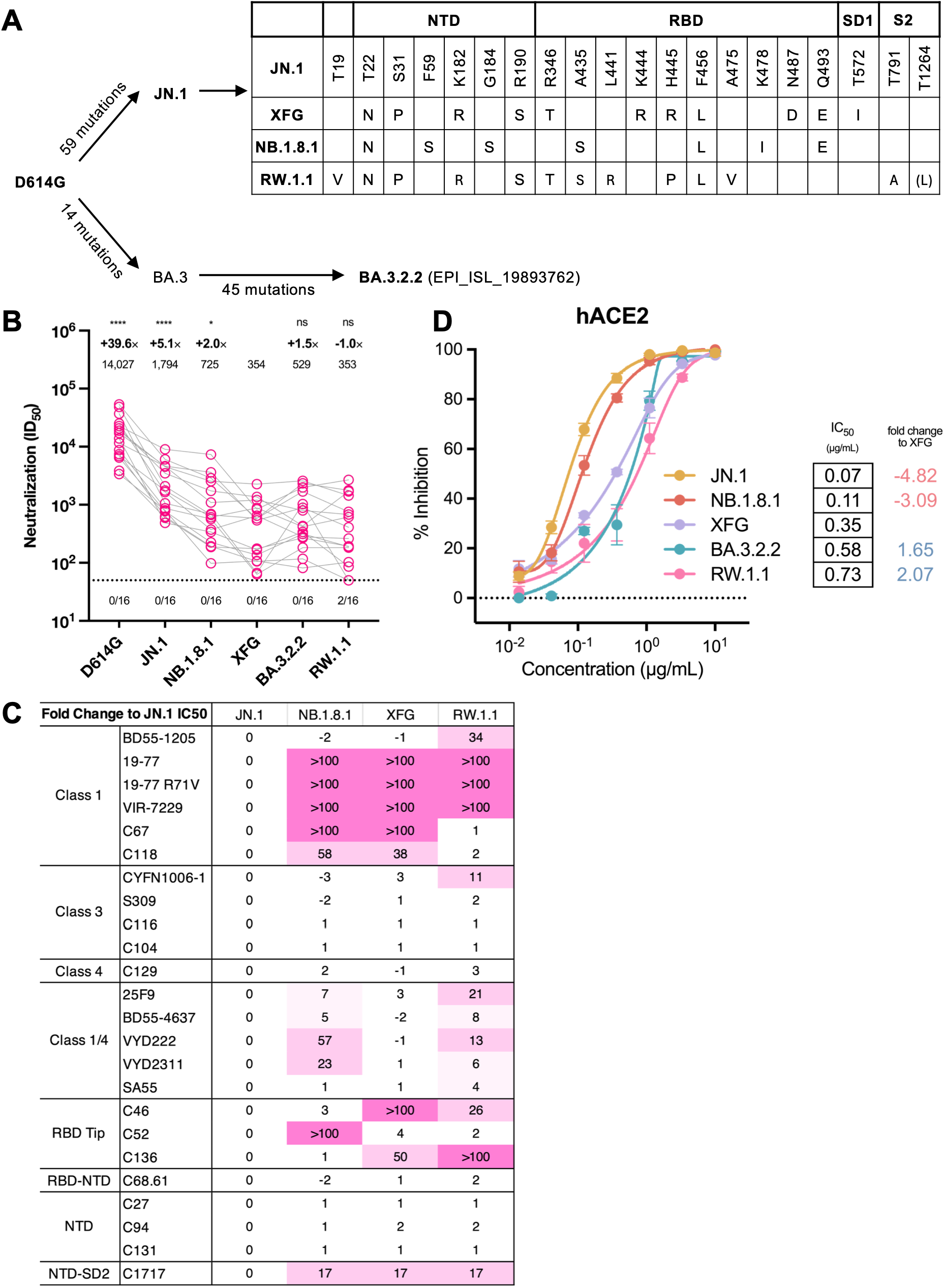
Antibody evasion and receptor binding of SARS-CoV-2 RW.1.1. A. Spike mutations present in selected SARS-CoV-2 variants, as tested here. Note that the Pango reference RW.1.1 also contains T478. B. Serum pseudovirus neutralization titers (ID_50_) against RW.1.1 and other variants. The geometric mean titer (GMT) is presented above each group. The fold change in GMT for each virus compared to XFG is also shown immediately above the GMT. Statistical analyses used Wilcoxon matched-pairs signed-rank tests, compared to XFG. ns, not significant. ∗: p < 0.05, ∗∗: p < 0.01, ∗∗∗: p < 0.001, ∗∗∗∗: p < 0.0001. Numbers under the dotted lines denote numbers of serum samples that were under the limit of detection (ID_50_ < 50). C. Fold change in neutralization IC_50_ values relative to JN.1 for the included mAbs, grouped by epitope classes. See **Table S1** for exact IC_50_ values. D. Sensitivity of 5 recent JN.1 subvariants to hACE2 inhibition. Data are shown as mean ± standard error of mean (SEM) for three technical replicates. Fold changes of variants are colored in red to indicate greater affinity to ACE2 than XFG or in blue to indicate lower affinity to ACE2 than XFG.

We first evaluated serum antibody evasion using pseudovirus neutralization assays with sera from 16 adults. Neutralizing titers against RW.1.1 were comparable to those against currently dominant variants, with a geometric mean titer (GMT) similar to that of XFG (dilution factor of 353 versus 354), which accounts for a plurality of infections in North America, and BA.3.2.2 (529), though lower than titers against NB.1.8.1 (725) (**Figure 1B**). Given our previous observation that BA.3.2.2 exhibited greater neutralization evasion in young children than in adults^3^, we additionally evaluated sera from infants and toddlers aged 6–28 months (n=12) and children aged 3–10 years (n=12). In contrast to BA.3.2.2^3^, neutralization of RW.1.1 did not differ significantly across age groups (**Figure S2**). Note that RW.1.1 spike tested herein bears K478, while the Pango reference RW.1.1 contains T478.

We next assessed whether RW.1.1 exhibited epitope-specific alterations in antibody recognition using a panel of 24 monoclonal antibodies targeting defined spike epitopes^4^. Despite showing no further increase in overall serum antibody evasion in the tested cohort, RW.1.1 exhibited increased evasion of several receptor-binding domain-directed (RBD) class 1 antibodies and increased evasion of some class 1/4 antibodies compared to previous strains (**Figure 1C, Table S1**).

Finally, we assessed spike-mediated receptor engagement using soluble hACE2 inhibition. RW.1.1 exhibited reduced hACE2 engagement compared with XFG, which itself has reduced receptor engagement relative to earlier JN.1 subvariants (**Figure 1D**). This further loss of receptor engagement is notable because XFG has outcompeted NB.1.8.1 despite reduced ACE2 affinity, again suggesting that higher receptor-binding affinity is not necessarily required for the epidemiologic expansion of recent SARS-CoV-2 variants.

Together, these findings suggest that RW.1.1 represents a continuation of the recent evolutionary trajectory exemplified by XFG, in which reduced receptor engagement is tolerated alongside changes in antibody evasion^2,5^. However, unlike the transition from LP.8.1.1 to XFG^2^, RW.1.1 does not exhibit a substantial further increase in overall serum antibody resistance, at least in the tested cohort. Instead, its antigenic evolution is characterized by selective evasion of specific antibody classes. Additional testing of potential effects of the K478T reversion substitution on antibody evasion and receptor binding are needed and ongoing.

As the composition of neutralizing antibody responses differs across populations according to vaccination and infection histories, selective escape from particular antibody specificities could potentially confer a growth advantage in subpopulations even in the absence of greater overall serum resistance in the tested cohort. The emergence of RW.1.1 therefore highlights how increasingly heterogeneous population immune landscapes may shape distinct evolutionary trajectories of SARS-CoV-2 and reinforces the need to evaluate not only overall serum neutralization but also the antibody specificities escaped by emerging variants.

## Notes

### Competing Interest Statement

D.D.H. co-founded TaiMed Biologics and RenBio, and he serves as a consultant for Brii Biosciences and is a board director at Vicarious Surgical. A.G. served as a member of the scientific advisory board for Janssen Pharmaceuticals and has consulted and serves on a scientific advisory board for Sanofi Pasteur.

### Summary of Updates

Text and Figure legend updated to clarify explanation of included mutations for one viral variant relative to reference database with a focus on spike amino acid position 478.

### Funding

- National Institutes of Health: subcontract no. 0258-A700-4609 under federal contract no. 75N93021C00014, subcontract GR0010139-PO024016 under federal contract no. 75N93021C00016, K08AI196255
- Columbia University: UR014016

