## Supplementary Appendix for "Antibody evasion and receptor binding of SARS-CoV-2 RW.1.1"

### Methods

#### Clinical Cohorts

Samples in the infants/toddlers and school-age children groups were procured through the Columbia University Center for Advanced Laboratory Medicine (CALM) in accordance with the protocol AAAO2000 approved by the Columbia University IRB. Residual whole blood samples were collected from children who had presented for lead testing by the clinical laboratories of New York-Presbyterian/Columbia University Irving Medical Center. Samples were selected and de-identified from children at least 6 months old and up to 10 years old who did not have a documented concurrent positive test result for SARS-CoV-2 by CALM. Samples in the adult group were collected in late 2025 before a vaccine booster dose through the VIVA study at the University of Michigan and through the "COVID-19 Persistence and Immunology Cohort (C-PIC)" study at Columbia University. Specimens were obtained following participant informed consent, adhering to the protocols approved by the IRBs of University of Michigan Medical School (protocol: HUM00232359) and Columbia University (protocol AAAS9722).

For pediatric samples: samples were centrifuged and plasma was aliquoted upon receipt. Samples were screened for SARS-CoV-2 viremia by RT-PCR and excluded if positive and screened for the presence of any anti-SARS-CoV-2 spike-binding antibodies using ELISA and included if any spike-binding antibodies were detected. All serum and plasma samples were heat inactivated at 56°C for 30 minutes before use.

Participants in the adult cohort were 81% female, 19% male, with an average age of 38.5 years. Participants in the school-age children cohort were 58% female, 42% male, with an average of 4.7 years. Participants in the infants/toddlers cohort were 50% female, 50% male, with an average age of 1.7 years. Details are summarized in **Table S2**. Although pediatric vaccination and infection history documentation was not accessible, the ELISA screening enabled more direct comparison against the adult cohort that did have a known prior COVID-19 vaccination and/or SARS-CoV-2 infection history. All adult participants had known vaccination histories, including but not limited to a minimum of three doses of mRNA vaccines targeting the ancestral strain of SARS-CoV-2. Individual adult vaccination histories are listed in **Table S3**.

#### Cell Lines

Vero-E6 (CRL-1586) cells and HEK293T (CRL-3216) cells were obtained from ATCC and cultured at 37°C with 5% $CO_2$ in Dulbecco's Modified Eagle Medium (DMEM) + 10% fetal bovine serum (FBS) + 1% penicillin-streptomycin. Vero-E6 cells are derived from African-green monkey kidneys. HEK293T cells and Expi293 cells are of human female origin. Expi293 (A14527) cells were purchased from Thermo Fisher Scientific and maintained in Expi293 medium per the manufacturer's instructions.

#### Plasmid Generation

As previously described, antibody sequences for the heavy chain variable (VH) and the light chain variable (VL) domains were synthesized by GenScript and then cloned into the gWiz vector to produce antibody expression plasmids. For the packaging plasmids for pseudoviruses, the spike constructs for D614G, JN.1, NB.1.8.1, XFG, and BA.3.2.2 were previously generated. The spike construct for RW.1.1 was designed as previous constructs and synthesized by Twist Biosciences. All constructs were verified using Sanger sequencing prior to use.

**Protein expression and purification**

The gWiz-antibody or pcDNA3-sACE2-WT(732)-IgG1 (Addgene 154104) plasmids were transfected into Expi293 cells using PEI-MAX at a ratio of 1:3. The supernatants were then collected after five days. The antibodies and human ACE2 (hACE2) fused to an Fc tag were purified with Protein A Sepharose (Cytiva) following the manufacturer's instructions. Prior to use, molecular weight and purity were confirmed by SDS-PAGE protein electrophoresis & size exclusion chromatography.

**Pseudovirus production**

VSV-based SARS-CoV-2 pseudoviruses were produced by replacing the native VSV glycoprotein with SARS-CoV-2 spike and its variants as previously described. Briefly, plasmids containing the appropriate spike were transfected into HEK293T cells with PEI Max. After 24 hours, VSV-G pseudotyped ΔG-luciferase (G*ΔG-luciferase, Kerafast) was added. The cells were washed with Dulbecco's Phosphate-Buffered Saline (DPBS) before being cultured in fresh medium for another 24 hours. Pseudoviruses were then harvested and centrifuged, then non-pseudotyped virus was eliminated using anti-VSVG (anti-I1) antibody. The resulting pseudovirus was then aliquoted and stored at -80°C.

**Pseudovirus neutralization assays with sera, mAbs, or ACE2**

Standardized viral infections dose was determined through SARS-CoV-2 pseudovirus titration. Seven serial dilutions of heat-inactivated sera, monoclonal antibodies (mAbs), and of soluble ACE2 were added in 96-well plates, starting at 1:50 dilution for sera, 10 µg/mL for antibodies, and 3 µg/mL for ACE2. As previously reported, we used soluble chimeric human ACE2 for ACE2 inhibition assays which contain ACE2 residues 1-732 fused to human IgG1 Fc. Pseudoviruses were then added and incubated at 37 °C for 1 hour. Control wells containing only pseudovirus were included in each plate. $4 \times 10^4$ Vero-E6 cells were then added to each well and incubated at 37 °C for 16 hours. Cells were lysed and luminescence was determined by the Luciferase Assay System (Promega E4550) and Tecan Infinite® 200 PRO using i-control™ software v.3.9.1.0, in accordance with the manufacturer's instructions.

**Quantification and statistical analysis**

Neutralization ID50 and IC50 values were determined by fitting a five-parameter dose-response curve in R (version 4.3.2) with the package drda (version 2.0.3).

### Author Contributions

The study was conceptualized by I.A.M. Experiments were conducted and data analyzed by K.D., H.H., C.Y.H., Y.G., and I.A.M. The results were analyzed, and the manuscript was written by K.D., H.H., Y.G., I.A.M., and D.D.H. The serum samples were collected or requested by A.G. and I.A.M. All contributing authors have reviewed and approved the manuscript.

### Acknowledgements

We thank Hiroshi Mohri (Columbia) for assistance with sample collection, Michael T. Yin, Jayesh G. Shah, Lawrence J. Purpura, Amanda Castillo, Meredith McNairy and Antonia Sturiza for conducting the C-PIC study (Columbia), and Carmen Gherasim, Virginia M. Pierce, Theresa Kowalski-Dobson, Anna Buswinka, Joseph Wendzinski, Mayurika Patel, Noah Paalanen and Ethan Hall of the VIVA study team for conducting the VIVA study. We thank Erin Poptanich, Tiffany Thomas, and Eldad Hod for advice and assistance related to requesting residual pediatric blood samples. We also thank all who share data on GISAID. This study was supported by funding from the NIH SARS-CoV-2 Assessment of Viral Evolution (SAVE) Program (subcontract no. 0258-A700-4609 under federal contract no. 75N93021C00014 to D.D.H and subcontract GR0010139-PO024016 under federal contract no. 75N93021C00016 to A.G.), K08AI196255 salary support to I.A.M., and internal startup funding UR014016 from Columbia University to Y.G.

### Declaration of Interests

D.D.H. co-founded TaiMed Biologics and RenBio, and he serves as a consultant for Brii Biosciences and is a board director at Vicarious Surgical. A.G. served as a member of the scientific advisory board for Janssen Pharmaceuticals and has consulted and serves on a scientific advisory board for Sanofi Pasteur.

### **Supplementary figures, tables, and legends**

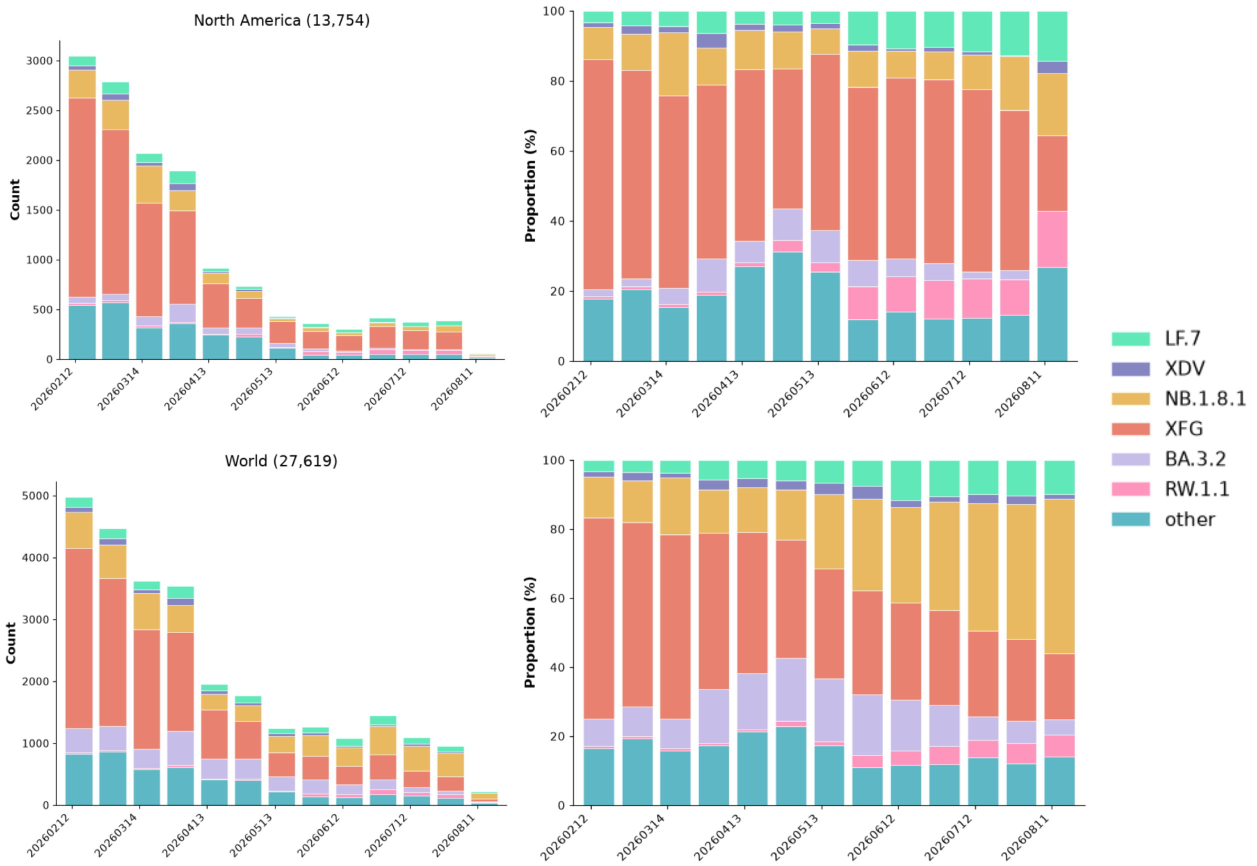

**Figure S1. Sequence frequency and relative frequencies of SARS-CoV-2 variants.**

Data were collected from February 12, 2026, to August 11, 2026, and stratified by region. Top panels show data from North America, and bottom panels show worldwide data. Left panels represent the sequency frequency, and right panels represent relative frequency of each variant. Numbers in parentheses indicate the number of sequences collected.

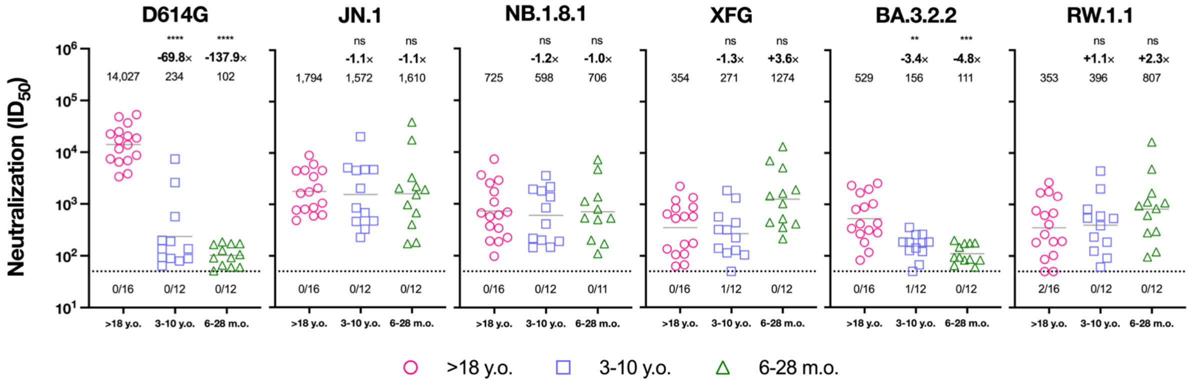

**Figure S2. Serum pseudovirus neutralization of SARS-CoV-2 RW.1.1 and other variants across age groups.**

Serum pseudovirus neutralization titers (ID50) against RW.1.1 and other variants. Note that one sample in the 6-28mo cohort tested against NB.1.8.1 was excluded due to low-quality luminescence readings.

| IC50 (μg/mL) | | D614G | BA.3.2.2 | JN.1 | NB.1.8.1 | XFG | RW.1.1 |
|---|---|---|---|---|---|---|---|
| Class 1 | BD55-1205 | <0.004 | 0.028 | 0.007 | <0.004 | 0.005 | 0.239 |
| | 19-77 | 0.014 | 0.580 | 0.095 | >20 | >20 | >20 |
| | 19-77 R71V | 0.016 | 0.033 | 0.009 | 2.225 | >20 | 6.545 |
| | VIR-7229 | 0.008 | 0.018 | 0.006 | 4.263 | 3.343 | 1.051 |
| | C67 | 19.582 | 6.430 | <0.004 | 1.051 | 5.050 | <0.004 |
| | C118 | >20 | >20 | <0.004 | 0.231 | 0.153 | 0.008 |
| Class 3 | CYFN1006-1 | <0.004 | 0.137 | 0.017 | 0.005 | 0.053 | 0.180 |
| | S309 | 0.041 | 12.302 | 9.881 | 4.639 | 12.115 | >20 |
| | C116 | >20 | 5.028 | <0.004 | <0.004 | <0.004 | <0.004 |
| | C104 | >20 | >20 | <0.004 | <0.004 | <0.004 | <0.004 |
| Class 4 | C129 | >20 | 1.889 | 0.328 | 0.812 | 0.293 | 1.053 |
| Class 1/4 | 25F9 | 0.011 | 11.502 | 0.970 | 7.164 | 2.465 | >20 |
| | BD55-4637 | 0.008 | 18.976 | 0.029 | 0.146 | 0.012 | 0.242 |
| | VYD222 | 0.007 | 1.970 | 0.253 | 14.528 | 0.231 | 3.299 |
| | VYD2311 | 0.012 | 0.112 | 0.008 | 0.187 | 0.010 | 0.050 |
| | SA55 | 0.010 | 0.008 | <0.004 | <0.004 | <0.004 | 0.017 |
| RBD Tip | C46 | >20 | >20 | 0.010 | 0.026 | >20 | 0.263 |
| | C52 | >20 | >20 | 0.006 | >20 | 0.024 | 0.013 |
| | C136 | >20 | >20 | <0.004 | <0.004 | 0.199 | >20 |
| RBD-NTD | C68.61 | 0.798 | 2.029 | 0.770 | 0.507 | 0.888 | 1.699 |
| NTD | C27 | >20 | >20 | <0.004 | <0.004 | <0.004 | <0.004 |
| | C94 | >20 | >20 | <0.004 | 0.005 | 0.007 | 0.006 |
| | C131 | >20 | >20 | <0.004 | <0.004 | <0.004 | <0.004 |
| NTD-SD2 | C1717 | 0.273 | 0.730 | 1.155 | >20 | >20 | >20 |

**Table S1. Neutralization $IC_{50}$s of included SARS-CoV-2 variants by mAb.**

Darker shades of red indicate lower $IC_{50}$.

| | All Participants | | Infants/Toddlers (0-28 m.o.) | | School-age children (3-10 y.o.) | | Adults (18+ y.o.) | |
|---|---|---|---|---|---|---|---|---|
| | No. (or Mean) | % (or Range) | No. (or Mean) | % (or Range) | No. (or Mean) | % (or Range) | No. (or Mean) | % (or Range) |
| Total | 40 | | 12 | | 12 | | 16 | |
| Female | 26 | 65% | 6 | 50% | 7 | 58% | 13 | 81% |
| Male | 14 | 35% | 6 | 50% | 5 | 42% | 3 | 19% |
| Age (y.o.) | 17.3 | (0,75) | 1.67 | (0,3) | 4.67 | (3,7) | 38.5 | (22,75) |

**Table S2. Cohort summary.**

y.o.: years old; m.o.: months old.

| ID | Age (Yr) | Sex | Race | No. Vax | No. WT Vax | No. BA.5 BV Vax | No XBB.1.5 MV Vax | No. KP.2 MV Vax | Sera Days Post Most Recent Infx Pre | Sera Days Post Most Recent Infx Post | Vaccine History |
|---|---|---|---|---|---|---|---|---|---|---|---|
| CUMC 1 | 26 | F | Asian | 6 | 3 | 1 | 1 | 1 | NA | NA | WT-P/WT-P/WT-P/BA.5-M/XBB.1.5-M/KP.2-M |
| CUMC 3 | 22 | M | Asian | 4 | 3 | 0 | 0 | 1 | 141 | 175 | WT-P/WT-P/WT-P/KP.2-P |
| CUMC 8 | 23 | F | Asian | 4 | 3 | 0 | 0 | 1 | NA | NA | WT-P/WT-P/WT-P/KP.2-M |
| CUMC 16 | 23 | M | Asian | 3 | 3 | 0 | 0 | 0 | 1030 | 1063 | WT-P/WT-P/WT-P |
| CUMC 18 | 23 | F | Asian | 3 | 3 | 0 | 0 | 0 | NA | NA | WT-S/WT-S/WT-S |
| MICH 1 | 48 | F | White | 4 | 3 | 0 | 0 | 0 | 896 | 927 | WT-P/WT-P/WT-P/XBB.1.5-P |
| MICH 2 | 30 | F | Black or African American | 5 | 3 | 1 | 0 | 1 | 1107 | 1148 | WT-P/WT-P/WT-P/BA.5-P/KP.2-P |
| MICH 3 | 31 | M | White | 5 | 3 | 0 | 1 | 1 | 901 | 953 | WT-P/WT-P/WT-P/XBB.1.5-P/KP.2-P |
| MICH 4 | 75 | F | More than One Race | 9 | 4 | 2 | 1 | 2 | NA | NA | WT-M/WT-M/WT-M/WT-M/BA.5-M/BA.5-M/XBB.1.5-M/KP.2-N/KP.2-P |
| MICH 5 | 33 | F | White | 6 | 3 | 1 | 1 | 1 | 260 | 286 | WT-P/WT-P/WT-P/BA.5-P/XBB.1.5-P/KP.2-M |
| MICH 6 | 26 | F | White | 4 | 3 | 1 | 0 | 0 | 1072 | 1115 | WT-P/WT-P/WT-P/BA.5-P |
| MICH 7 | 52 | F | White | 6 | 3 | 1 | 1 | 1 | NA | NA | WT-P/WT-P/WT-P/BA.5-P/XBB.1.5-M/KP.2-M |
| MICH 8 | 55 | F | White | 7 | 4 | 1 | 1 | 1 | 742 | 778 | WT-P/WT-P/WT-P/WT-P/BA.5-P/XBB.1.5-P/KP.2-N |
| MICH 9 | 29 | F | White | 5 | 3 | 1 | 0 | 1 | 430 | 473 | WT-P/WT-P/WT-P/BA.5-U/KP.2-U |
| MICH 10 | 60 | F | White | 6 | 3 | 1 | 1 | 1 | 1235 | 1276 | WT-M/WT-M/WT-M/BA.5-M/XBB.1.5-M/KP.2-M |
| MICH 11 | 60 | F | White | 5 | 3 | 1 | 0 | 1 | 1177 | 1220 | WT-M/WT-M/WT-M/BA.5-P/KP.2-M |

**Table S3. Adult participant vaccination histories.**

Vaccine formulations are denoted as wildtype (WT), BA.5 Bivalent (BA.5), XBB.1.5 monovalent (XBB.1.5), and KP.2 monovalent (KP.2). Vaccine manufacturers are denoted as Pfizer (P) or Moderna (M). Yr, years; Infx, infection; Vax, vaccination; F, female; M, male.
